# Genomic Considerations for FHIR; eMERGE Implementation Lessons

**DOI:** 10.1101/2021.01.31.429037

**Authors:** Mullai Murugan, Lawrence J. Babb, Casey Overby Taylor, Luke V. Rasmussen, Robert R. Freimuth, Eric Venner, Fei Yan, Victoria Yi, Stephen J. Granite, Hana Zouk, Samuel J. Aronson, Kevin Power, Alex Fedotov, David R. Crosslin, David Fasel, Gail P. Jarvik, Hakon Hakonarson, Hana Bangash, Iftikhar J. Kullo, John J. Connolly, Jordan G. Nestor, Pedro J. Caraballo, WeiQi Wei, Ken Wiley, Heidi L. Rehm, Richard A. Gibbs

## Abstract

Structured representation of clinical genetic results is necessary for advancing precision medicine. The Electronic Medical Records and Genomics (eMERGE) Network’s Phase III program initially used a commercially developed XML message format for standardized and structured representation of genetic results for electronic health record (EHR) integration. In a desire to move towards a standard representation, the network created a new standardized format based upon Health Level Seven Fast Healthcare Interoperability Resources (HL7 FHIR), to represent clinical genomics results. These new standards improve the utility of HL7 FHIR as an international healthcare interoperability standard for management of genetic data from patients. This work advances the establishment of standards that are being designed for broad adoption in the current health information technology landscape.

## 1. Introduction

The use of genomic testing in healthcare has grown in recent years, attributed to our increased understanding of the human genome and broader accessibility of tests[1–4]. Typically, the results from genetic testing laboratories are represented in a narrative, unstructured form, delivered as a Portable Document Format (PDF) document, which is simply scanned or uploaded to the electronic health record (EHR)[5]. This approach, though widely accepted and ubiquitous in its use, is primarily targeted to human readers. PDF and other custom structured reporting, though serving the immediate need of delivering genomic test results, hinders widespread reanalysis, data sharing, interoperability, automation, and downstream research efforts. Furthermore, without computational standards we are unable to reliably exchange genomic data and use those data to scale discovery and improve clinical care through automated clinical decision support (CDS)[6]. Because access to relevant genomic data in a standardized computable format has been a major hurdle to widespread implementation of genomic medicine, standards organizations, research networks and healthcare institutions have embarked on multiple projects in the past decade to improve the accessibility of genomic test results as computable data[7–9].

The Electronic Medical Records and Genomics (eMERGE) Network, funded by the National Human Genome Research Institute (NHGRI), is a consortium of U.S. medical research institutions that “develops, disseminates, and applies approaches to research that combine biorepositories with electronic medical record systems for genomic discovery and genomic medicine implementation research”[10]. Phase III of this program, as part of the Network’s overarching goal to integrate genomic test results to the EHR for clinical care, sought to create a proof of concept, standards-based representation of clinical genomic test results using the rapidly evolving Health Level Seven (HL7) Fast Healthcare Interoperability Resources (FHIR)[11] standard.

### 1.1. eMERGE Background

Currently in its fourth phase, the eMERGE Network’s first phase began in September 2007. eMERGE Phase III (September 2015 to March 2020) joined together multiple laboratories and clinics, including two central sequencing and genotyping facilities (CSGs): Baylor College of Medicine Human Genome Sequencing Center (BCM-HGSC) and the Broad Institute (BI) and Partners Laboratory for Molecular Medicine (LMM), as well as eleven study sites. The CSGs performed sequencing, variant interpretation, and clinical report generation. The reports were then returned to clinical sites and integrated into the respective EHRs[12,13]. As illustrated in Figure 1[14], the complexity and heterogeneity of Phase III of the Network provided considerable challenges for technical and data harmonization across the CSGs and study sites[15]. The network identified workflow and logistic components to harmonize both data and delivery of results, as well as the need for both a narrative PDF form to satisfy clinicians and regulatory requirements and a standardized XML format to represent clinical genetic test results directly in the EHR.

**Figure 1.**
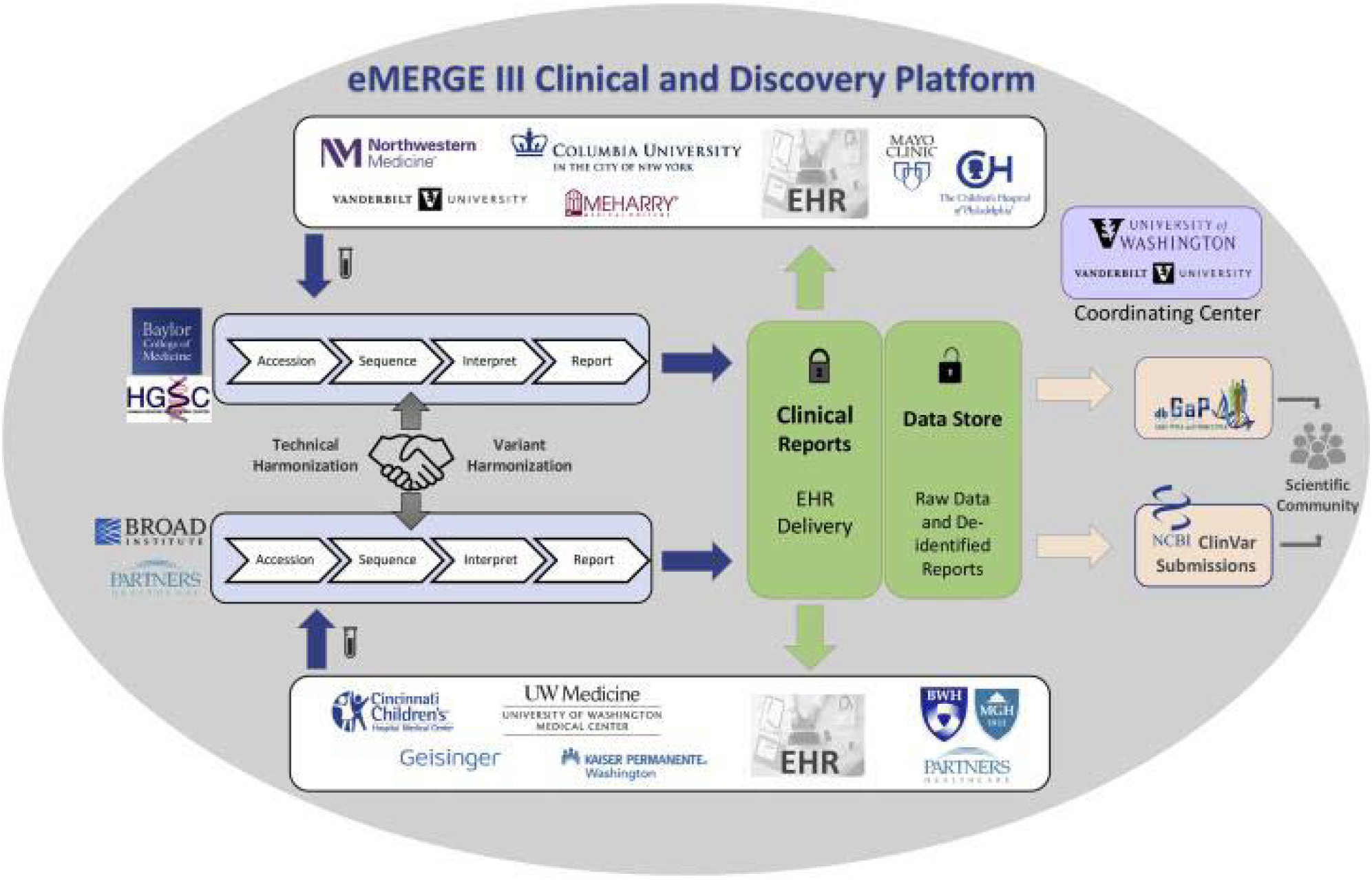
The eMERGE III Network. The Network comprised 11 study sites, 2 central sequencing and genotyping facilities (CSGs) and a coordinating center (CC) with samples sent from the study sites to the CSGs. Clinical Genetic Results were returned to the study sites after sequencing, variant classification and harmonization at the CSGs with raw data and de-identified results being returned to the CC for research. Both the clinical and research data flow is illustrated in this figure[14].

The electronic structure utilized by the eMERGE Network for genetic test reports included an XML format based primarily upon the features of Sunquest Mitogen^™^ Genetics (formerly known as GeneInsight), a commercial tool for clinical genetic test reporting and knowledge management originally developed by one of the CSGs[15]. This XML format enabled report data exchange between the eMERGE sites that used GeneInsight, the sites that employed local data, and the centralized repositories. The GeneInsight format, though opportune for the eMERGE Network at delivering the initial requirements, is not consistent with other open standards and thus not viable as the long-term solution required to support broader growth and adoption. With the emergence of FHIR as an interoperable healthcare standard and the work of the HL7 Clinical Genomics Workgroup (CG WG) towards the creation of a FHIR Genomics Reporting Implementation Guide (GR IG)[16], the Network decided on a specification based on the GR IG in an effort to contribute to and validate the nascent GR IG.

### 1.2. FHIR Background

The FHIR standard builds on the widely used HL7 Version 2[17] messaging standard and leverages common, modern web technologies that facilitate the development and integration of applications into clinical environments. FHIR has gained traction since its inception in 2012 as an open API standard for interoperability, continuing to grow with programs such as the *FHIR Accelerator Program*[18], the healthcare pledge by technology companies to adopt emerging standards such as FHIR for interoperability[19], along with NIH notices encouraging the use of FHIR for clinical and research interoperability[20,21]. Clinical software vendors are also increasingly implementing support for FHIR-based APIs into EHRs and laboratory information management systems (LIMS), enabling clinical lab test results to be rendered as discrete data elements that can be used to drive CDS.

Building upon the core FHIR standard, the FHIR community has worked to support its implementation for clinical genetics and genomics. In November 2019, the HL7 CG WG published the first release of the GR IG, which is intended to support “all aspects[22]” of reporting clinical genomics data. The GR IG contains specific guidance for reporting genomic variants that pertain to pharmacogenomics (PGx), tumor testing, and histocompatibility profiling. The current release of the GR IG [16], based on FHIR R4[22], has been balloted as a Standard for Trial Use (STU), which is intended to be vetted through implementation and pilot testing by early adopters. This process is critically important to the advancement of the standard as feedback drives improvements in the specification before it becomes more widely adopted, which limits the ability to make breaking changes as gaps are filled.

### 1.3. Project Objectives

In 2019 the eMERGE Network aimed to improve structured data standards and take advantage of newly emerging FHIR capabilities. Three milestones were established: 1. Develop a computable and standardized clinical reporting specification for eMERGE with FHIR; 2. Create a proof-of-concept implementation pilot, generating eMERGE Clinical Genetic Reports with the eMERGE FHIR Specification; and 3. Establish EHR interoperability with FHIR-enabled secure ingestion of genetic results and implement clinical decision support use cases.

Recognizing the emerging nature of this landscape, the network limited the scope of its effort to building a baseline upon which future work can be continued. Here we describe these efforts, focusing on identifying lessons learned and future considerations on how organizations can begin to adopt the FHIR genomics standard, illustrating the level of expertise and effort required to do so, and demonstrating the positive impact that such efforts can have on the development of international standards.

## 2. Methods

Fulfillment of the objectives and the milestones set forth for this project was managed as a cross-network collaboration with the two CSGs first developing the eMERGE FHIR Specification. The BCM-HGSC acted in its role of a diagnostic laboratory to generate a sample set of clinical genetic reports utilizing the specification, and study site Northwestern University (NU) and non-clinical affiliate Johns Hopkins University (JHU) acted as provider facilities that implemented a proof-of-concept EHR integration and clinical care pilot to demonstrate the viability of the eMERGE FHIR Specification. These steps are illustrated in Figure 2 and detailed further in this section.

**Figure 2.**
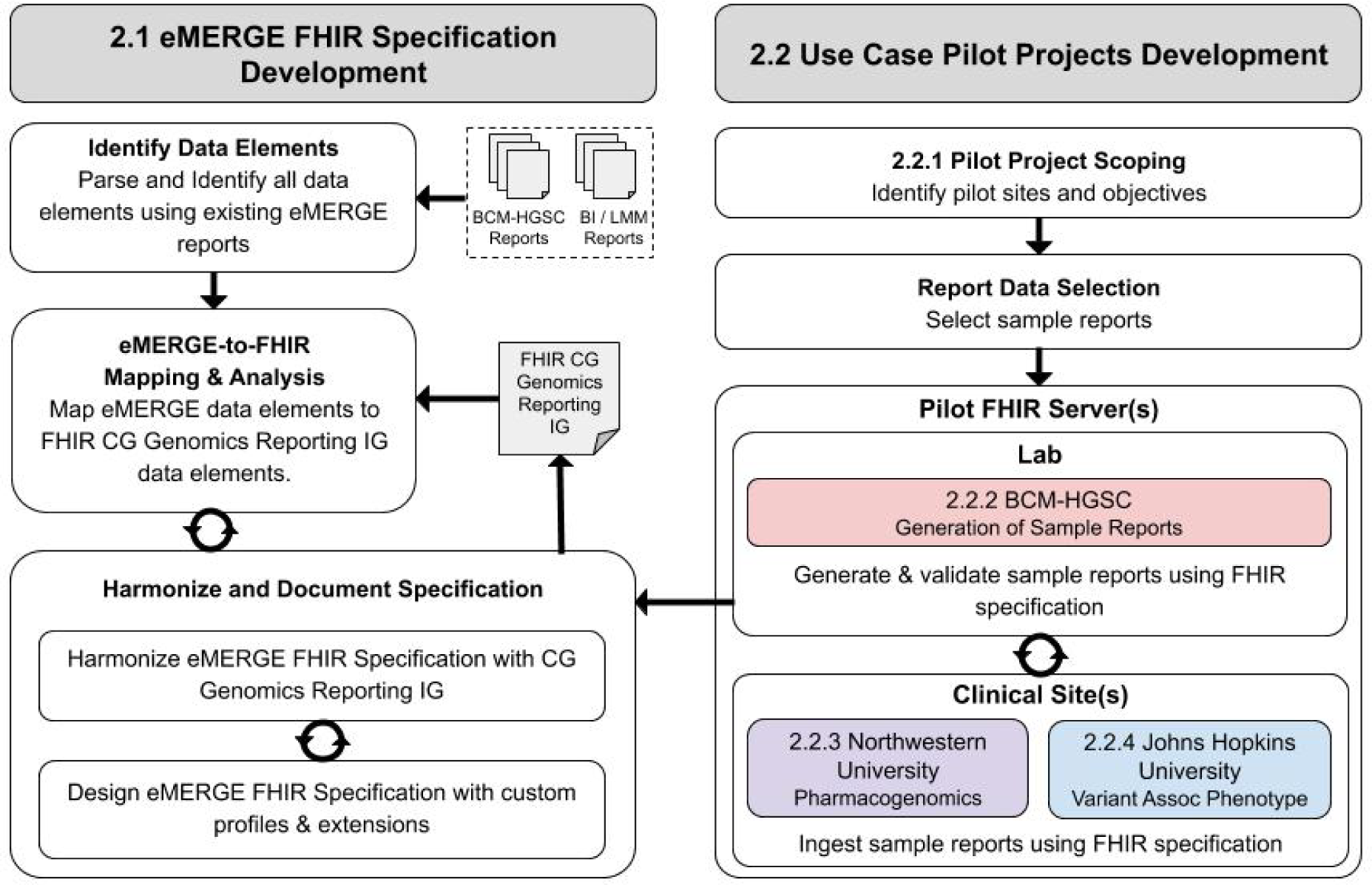
Methods for specification and pilot projects development. The two columns represent complementary work streams - the leftmost dedicated to the specification development, and the right on use cases for pilot projects development. The solid arrows represent the stepwise or iterative direction of the respective processes.

### 2.1. eMERGE FHIR Specification Development

The two CSGs established the following guiding principles for this project, taking advantage of their respective experiences:

- **Structured content -** All content from the narrative PDF eMERGE reports and all eMERGE standard reporting use cases should be captured in structured format and as meaningful data elements without losing content and context;
- **Alignment with HL7 FHIR Core and GR IG** - All eMERGE concepts and associated elements shall be aligned with GR IG and FHIR Core Standards and extended as required;
- **Computationally reliable representation of results** - An optimal computational form for each data element shall be determined, prioritizing eMERGE pilot objectives for HER integration and CDS. All specimen types and genetic data elements related to the resulting observations must be based on reference sequences, coordinates and structures that consistently and accurately reflect the lab methods used to align the raw data, determine coverage and call the variants; and
- **Codify concepts when reasonable** - Concepts should be codified using FHIR Core and GR IG guidance. eMERGE concepts that extend beyond the FHIR and CG guidance should be codified if possible and within reason.

The development of the eMERGE FHIR Specification consisted of the following steps -

- **Identify data elements** using existing results from a comprehensive set of eMERGE reporting use cases;
- **Map eMERGE report** elements and structures both semantically and structurally to GR IG resources and profiles; perform an analysis, identify issues that require further resolution, and propose resolutions; and
- **Harmonize and document** finalized decisions informed by
  - Harmonizing changes with GR IG
  - Documenting resolutions requiring custom profiles & extensions
  - Feedback from BCM-HGSC lab pilot development

The steps defined above were accomplished by first analyzing eMERGE narrative reports representing a range of use cases. Next, working with domain experts (geneticists from both the CSGs), the semantic meaning and structure of these concepts were verified and all data elements appearing on the reports were associated with the proper concepts. This step was critical to assuring that at a minimum both CSGs agreed with the conceptual meaning of all granular elements on their reports, minimizing subsequent misrepresentation of the elements.

The next step was to determine how best to computationally represent these concepts. A gap analysis was performed between each eMERGE concept and its corresponding GR IG resources and profiles. The aim of this effort was to synchronize GR IG resource profiles to related eMERGE reporting concepts. Gaps, questions and issues arising during this process were publicly documented and harmonized between the eMERGE CSGs and the HL7 FHIR community. This communication consisted of informal feedback through the HL7 FHIR Zulip Chat Board[23], and formal feedback through issue trackers on Jira[24]. Complex questions were addressed through discussions with the Clinical Genomics WG or the eMERGE EHRI Workgroup. This reconciliation and harmonization process resulted in a beta version of the eMERGE FHIR Specification. This process also contributed to enhancements and changes to the GR IG itself. Implementation of the use case pilot projects contributed to the validation of the eMERGE FHIR Specification and generated additional feedback that was integrated into the specification.

### 2.2. Use Case Pilot Projects Development

#### 2.2.1. Pilot Project Scoping

We conducted a proof-of-concept implementation pilot that represented real clinical genetic testing reports using the eMERGE FHIR Specification in EHRs. The pilot project included three use cases: 1. BCM-HGSC - Generation of sample reports using the eMERGE FHIR Specification, 2. NU - Pharmacogenomics clinical decision support, and 3. JHU - Variant associated phenotype. The pilot further allowed ratification of the eMERGE FHIR Specification via iterative feedback from the test sites.

#### 2.2.2. BCM-HGSC - Generation of Sample Reports

The pilot included representative data from a wide cross section of samples, including positive and negative results, heterogeneous indications for testing, multiple variant and PGx findings, selected phenotype susceptibilities, sex, and race. Reports for the sample cohort were generated in the eMERGE FHIR Specification.

The BCM-HGSC developed a custom FHIR Client[25] using the HAPI FHIR API[26] to convert the raw JSON results output of the BCM-HGSC’s Clinical Pipeline Neptune to FHIR Bundles consisting of resources, profiles, extensions and coding systems as identified in the specification. A secure HIPAA compliant Amazon Web Services (AWS) cloud-based SMILE CDR[27] environment was established with the necessary GR IG profiles, FHIR extensions, and relevant coding systems such as Logical Observation Identifiers Names and Codes (LOINC), Systematized Nomenclature of Medicine (SNOMED), and the HUGO Gene Nomenclature Committee (HGNC). This environment was used to validate the structure and content of the FHIR Bundles. The FHIR Bundles themselves were stored on the SMILE CDR server and on AWS S3.

During this implementation, the group iteratively revised and refined the specification based on the implementation’s efficacy assessment, and validated results with FHIR Core and GR IG to ensure adherence. Finally, the BCM-HGSC tested and internally validated FHIR bundles, representing the sample cohort, for data integrity and accuracy using the narrative PDF. The content of the FHIR reports such as patient/sample identifiers and results were validated against the source JSON via automation. A set of three FHIR reports were manually reviewed end to end for structure, format and content by the developer (F.Y.). Once validated, FHIR bundles were distributed to JHU and NU.

#### 2.2.3. Northwestern University - Pharmacogenomics

The NU use case targeted integrating FHIR results into an existing Ancillary Genomics System[28] (AGS), which was used as an intermediary platform to transmit results to the EHR. Pre-existing PGx CDS alerts were configured within the EHR to alert providers to recommendations for clopidogrel, warfarin, and simvastatin, and are available from the CDS Knowledgebase[29].

In order to complete this use case, NU required access to the PGx interpretations within the overall report, including genes, diplotypes, and medications. Because of institutional constraints surrounding the use case, the system was not connected to the live EHR environment. Instead, given that a validated workflow existed for the same data, we considered success as the ability to import the FHIR bundles into a test FHIR server, and the ability to transform the FHIR resources into the same intermediate results that were sent to the EHR to trigger CDS.

NU implemented a locally hosted instance of the open-source Microsoft FHIR Server[30], connected to a Microsoft SQL Server 2017 Docker container as the underlying data store, and configured with the FHIR R4 Specification. A command line application was created in C# using the .NET Core framework, and is freely available on GitHub[31]. This utility loaded FHIR bundles stored on disk to the FHIR server in order to simulate the process of “receiving” results from an external laboratory. In addition, the utility then extracted data from the FHIR Server to the simulated AGS. The import process was manually validated for a set of 25 randomly selected patients by one of the authors (L.V.R) to confirm the correct attribution of diplotypes to patients.

#### 2.2.4. Johns Hopkins University - Variant Associated Phenotype

The goal of the JHU use case was to document in the EHR the phenotype(s) known to be associated with a pathogenic or likely pathogenic variant and the corresponding evidence provided by the diagnostic lab in the FHIR report. A cloud based FHIR service for FHIR R4 and DSTU3 was deployed as an AGS to provide EHR access to genomic lab report data. In addition, a secure FHIR Genomics Proxy instance was deployed[32] with an authorization access role for BCM-HGSC users to send FHIR bundles to the AGS. With proper authentication, JHU accepted FHIR-formatted reports from BCM-HGSC and stored those reports on the cloud-based FHIR server. JHU used the Azure API for FHIR[33] and registered it in the JHU API portal[34] so that it could be used to add a “variant associated phenotype” to the patient chart for a set of test patients using a web service supported by Epic(R)[35].

In order to validate the JHU import process, a BCM-HGSC application was generated to create FHIR transaction bundles and submit them to the JHU FHIR server, while the JHU team generated a Validator tool. The number of bundles submitted by BCM-HGSC and OMIM identifiers in those reports found in the JHU AGS were reported. (See **Appendix A** Johns Hopkins University Technical Validation).

#### 2.2.5. Assessment of Data Quality for Use Cases

For each use case we assessed data quality characteristics of the electronic patient data used by the NU and JHU sites for their CDS use cases. Our approach adapted a data quality scoring system to score the FHIR transaction bundles according to four characteristics (Completeness, Accessibility, Relevance/Fitness, and Reliability/Accuracy). **Appendix B** provides details about the methods to assess data quality that NU and JHU applied after completing their use cases.

## 3. Results

### 3.1. eMERGE FHIR Specification Development

The principal outcomes of the specification development were the following: 1. To identify the complete set of report concepts and elements used throughout all eMERGE reporting use cases; 2. To create a FHIR based schema using the GR IG that would be implementable by eMERGE; 3. To provide a public document of the eMERGE FHIR Specification; and 4. To collaborate with the HL7 CG WG to harmonize useful feedback into the GR IG and FHIR Specification in general.

#### 3.1.1. Identification of eMERGE Report Concepts and Elements

The first step towards the creation of the eMERGE FHIR Specification was an “As Is” analysis of the existing genetic reports to inventory all eMERGE reporting concepts and elements. To this end, we compiled a set of all-inclusive representative reports from both the CSGs (see Figure 3 for an example report from each CSG) to ensure use cases requiring unique report concepts and elements were included. Table 1 contains the list of principal eMERGE reporting scenarios.

**Table 1:**
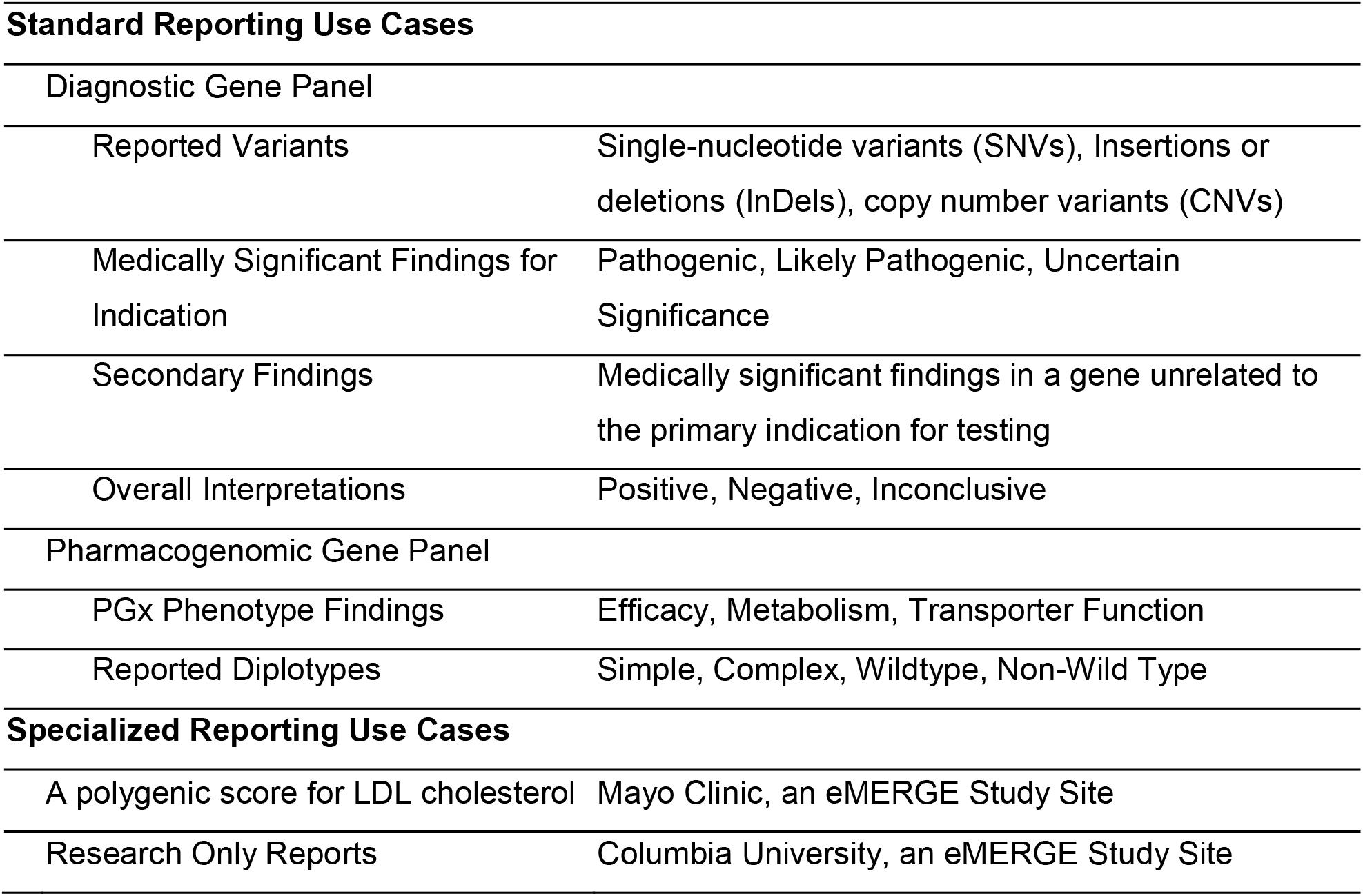
eMERGE Reporting Use Cases. This table lists all of the eMERGE reporting use cases. The first column is a breakdown of reporting result types and the second column lists more specific outcomes for the corresponding reporting result type. The specialized reporting use cases are specified for the associated eMERGE study sites.

**Figure 3.**
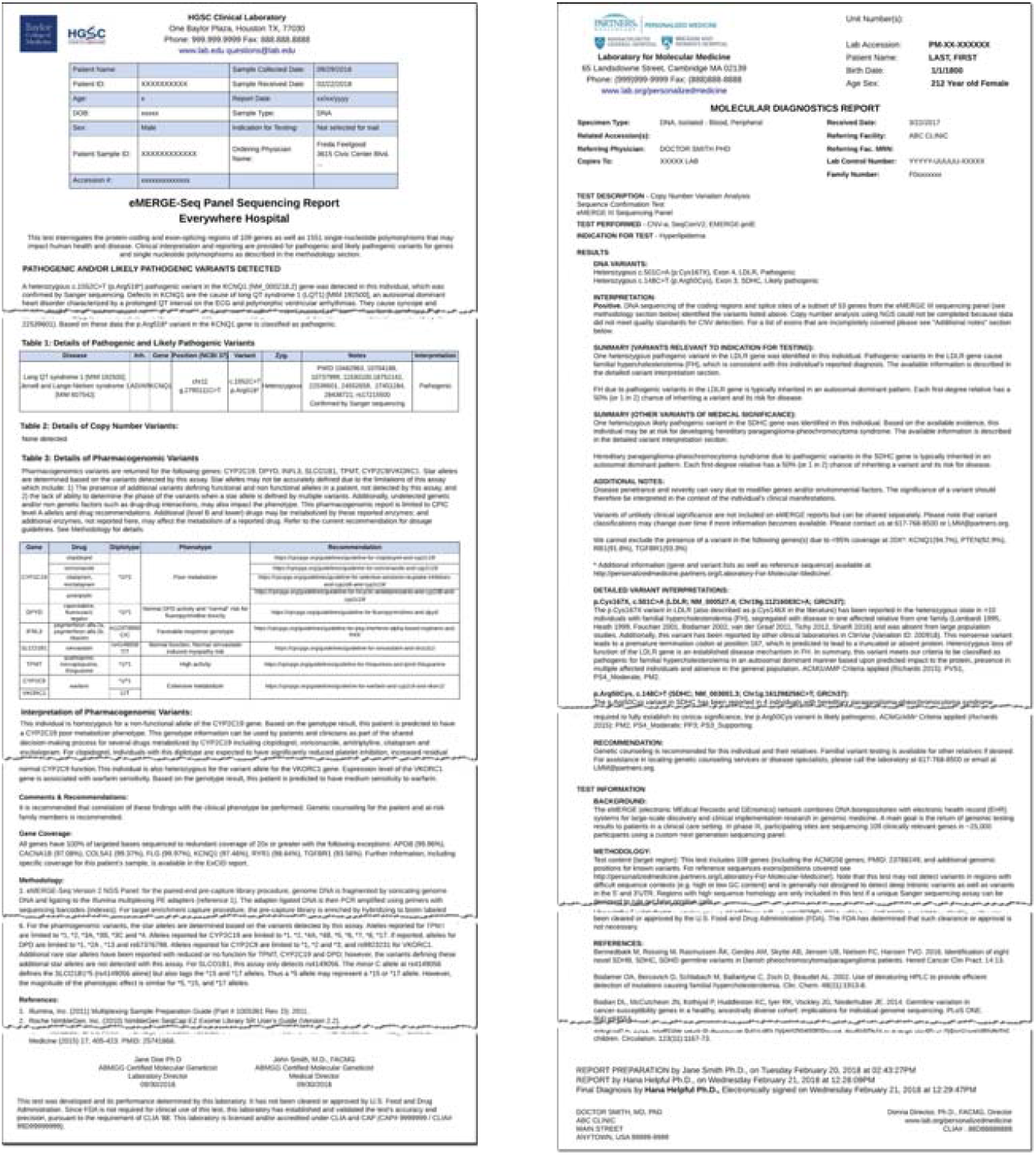
BCM-HGSC & LMM eMERGE Narrative Report Examples. Reports illustrate the content and structure of a typical positive genetic report generated by each of the CSGs.

Using selected reports for these associated use cases, the structure and composition of the reports was analyzed, and a set of data elements was assembled (Figure 4), resulting in 18 core concepts and around 100 fundamental data elements. This analysis and documentation of the existing eMERGE report content served as the foundation for the design of the eMERGE FHIR Specification.

**Figure 4.**
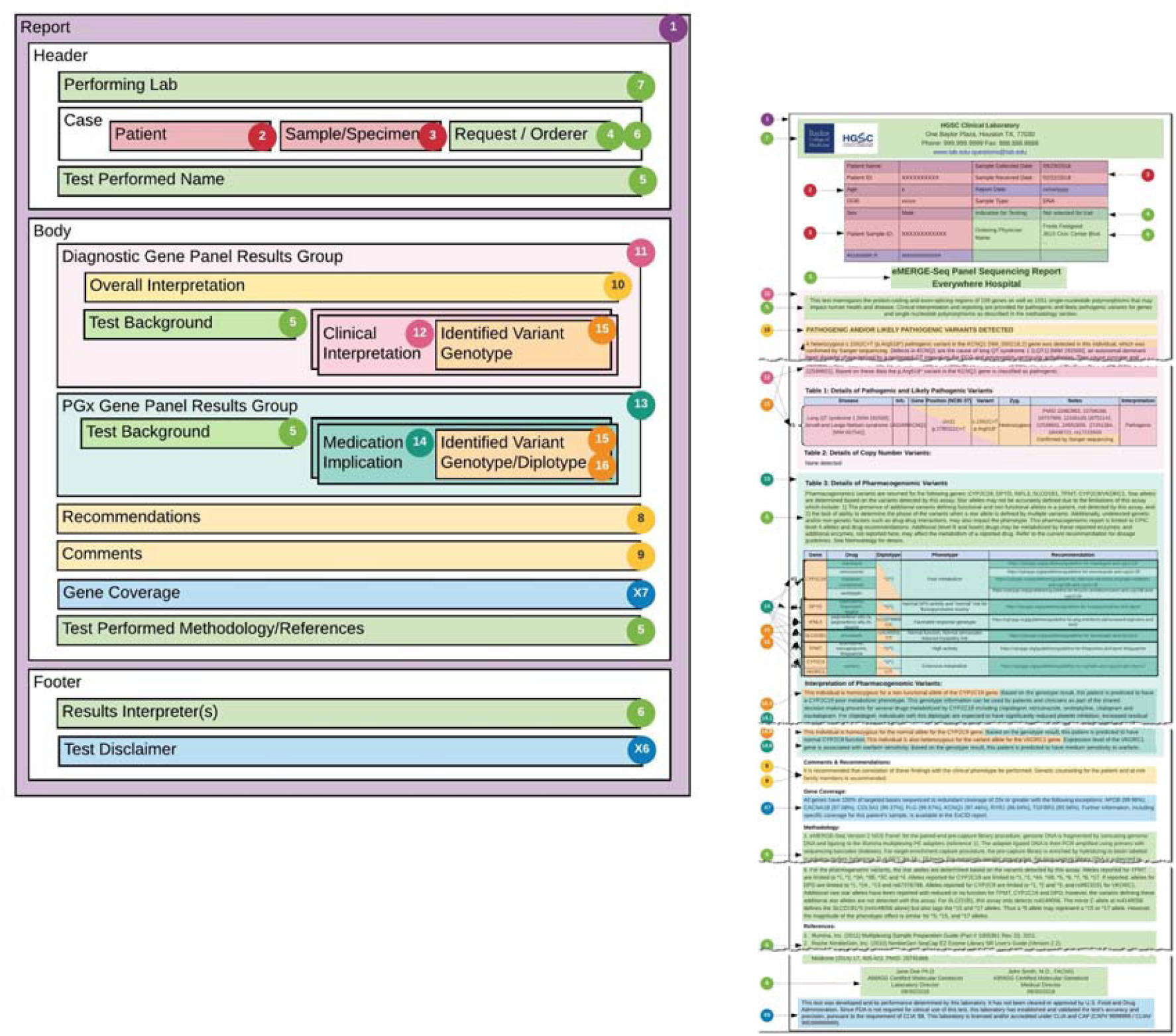
BCM-HGSC general report layout and detailed mapping. Illustrates the composition and layout of the data concepts within the eMERGE reports with each of the concepts coded by color and more precisely by number. Note: though Figure 4 is the BCM-HGSC format, BI/LMM also included the same concepts in a slightly modified layout. The numbering also correlates to the concepts in Figure 5.

In an effort to manage the pilot within allotted resources and time, the scope of this effort was confined to the Standard Reporting Use Cases and result delivery, while including provisions for future expansion.

#### 3.1.2. eMERGE Report to FHIR GR IG - Mapping and Analysis

The next step in the development of the eMERGE FHIR Specification was the mapping of eMERGE report concepts and elements to the GR IG. Adopting the GR IG’s guidance[16], all major eMERGE report concepts were aligned to the GR IG resources and profiles, followed by a granular mapping of every eMERGE report element to a corresponding FHIR resource element.

##### 3.1.2.1. Mapping eMERGE Report Concepts to FHIR Resources and Profiles

The GR IG provided the guidance for driving the mapping of the eMERGE report concepts to its resources, profiles and extensions. Our first attempt at mapping resulted in several key structural and organizational questions; as these questions and issues were fundamental to the design of the eMERGE FHIR Specification, a significant amount of time was spent to identify long and short-term resolutions.

The end result was a set of 21 noteworthy issues and associated resolutions, documented as part of the eMERGE Specification[36]. Below is a representative issue and its resolution followed by a complete listing of all 21 issues that were addressed (Table 2). These issues and their resolutions vary in scope and complexity with proposed solutions being driven by balancing the timeline for delivering the eMERGE pilot against the ideal solution as noted.

**Table 2:**
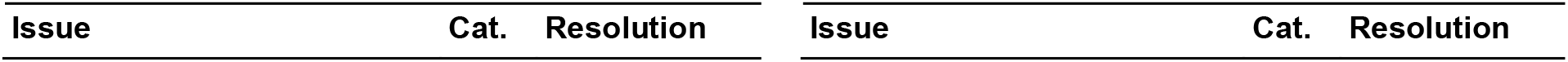
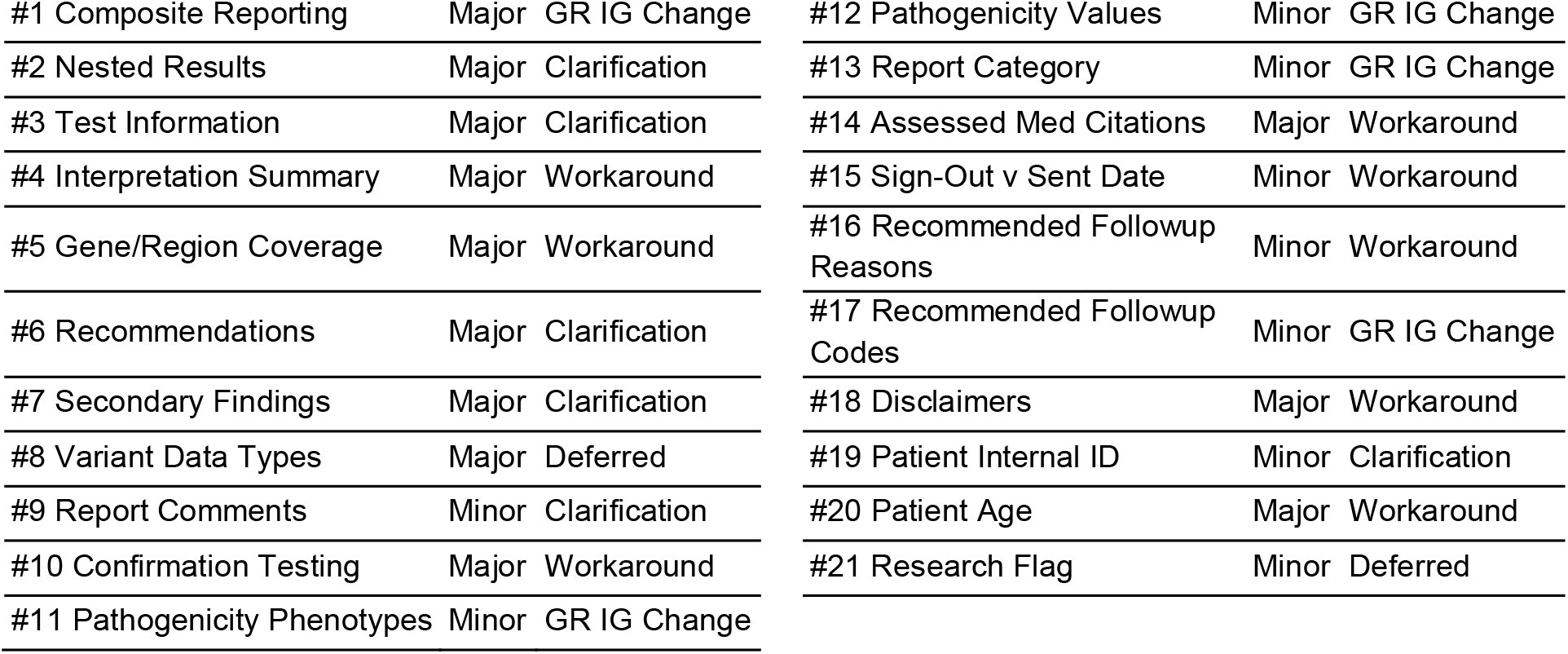
Issues & Resolutions. This table lists all of the noteworthy issues that required investigation and collaboration with the HL7 CG WG to resolve. The three columns show the issue label, category (*Major, Minor*) and resolution status (*Clarification* - GR IG or FHIR solution, *STU1 Change* - CG WG modified GR IG to support, *Workaround* - extension or non-standard approach, *Deferred* - deferred workaround but submitted for future consideration). Detailed description for each of these issues is available on the Issues & Resolutions[36] page of the eMERGE Read the Docs website[40].

**#1 Composite reporting[37]**

Diagnostic labs frequently include several types of results in genetic test reports which then appear in separate sections within an overarching report. eMERGE genetic reports are examples of this model and include both gene panel interpretation as well as PGx results. This style of reporting is analogous to the notion of composite reporting whereby while individual sections of the report can be treated independently, they are still combined together as they rely on shared findings from an upstream wet lab assay such as Whole Exome Sequencing or Gene Panels. We evaluated two options to represent composite reporting: 1. Nested Diagnostic Report Resources or 2. The Observation Grouper Profile[38]. Based on analysis and collaborative discussions with the CG WG[39], we decided on the Grouper Profile.

The resolution to use the Observation Grouper Profile allows consuming EHR systems to utilize the Gene Panel & PGx results as disparate components. Furthermore, while the usage of Grouper Profile was well-defined by the GR IG, the concept of nested Diagnostic Reports was still under investigation and not ready for adoption. The benefit of this approach also lends itself to including additional sections such as Polygenic Risk Scores to genetic reports.

Addressing and resolving these issues resulted in the mapping and structural design of the specification, illustrated in Figure 5. As illustrated, the root profile of the specification is the GenomicsReport; this is the key resource that encapsulates the ServiceRequest for the test, the Observations that constitute the results (i.e. findings or implications of the test), the Tasks that include clinical care recommendations, and the Grouper Profile to organize and manage composite resulting (i.e. GenePanel and PGx results). Other major resources attached to the GenomicsReport include the Patient for whom the test is being ordered, the associated Specimen, the Practitioner ordering the test, the Organization (i.e. Diagnostic Laboratory performing the test) and the Practitioner interpreting the results of the test.

**Figure 5.**
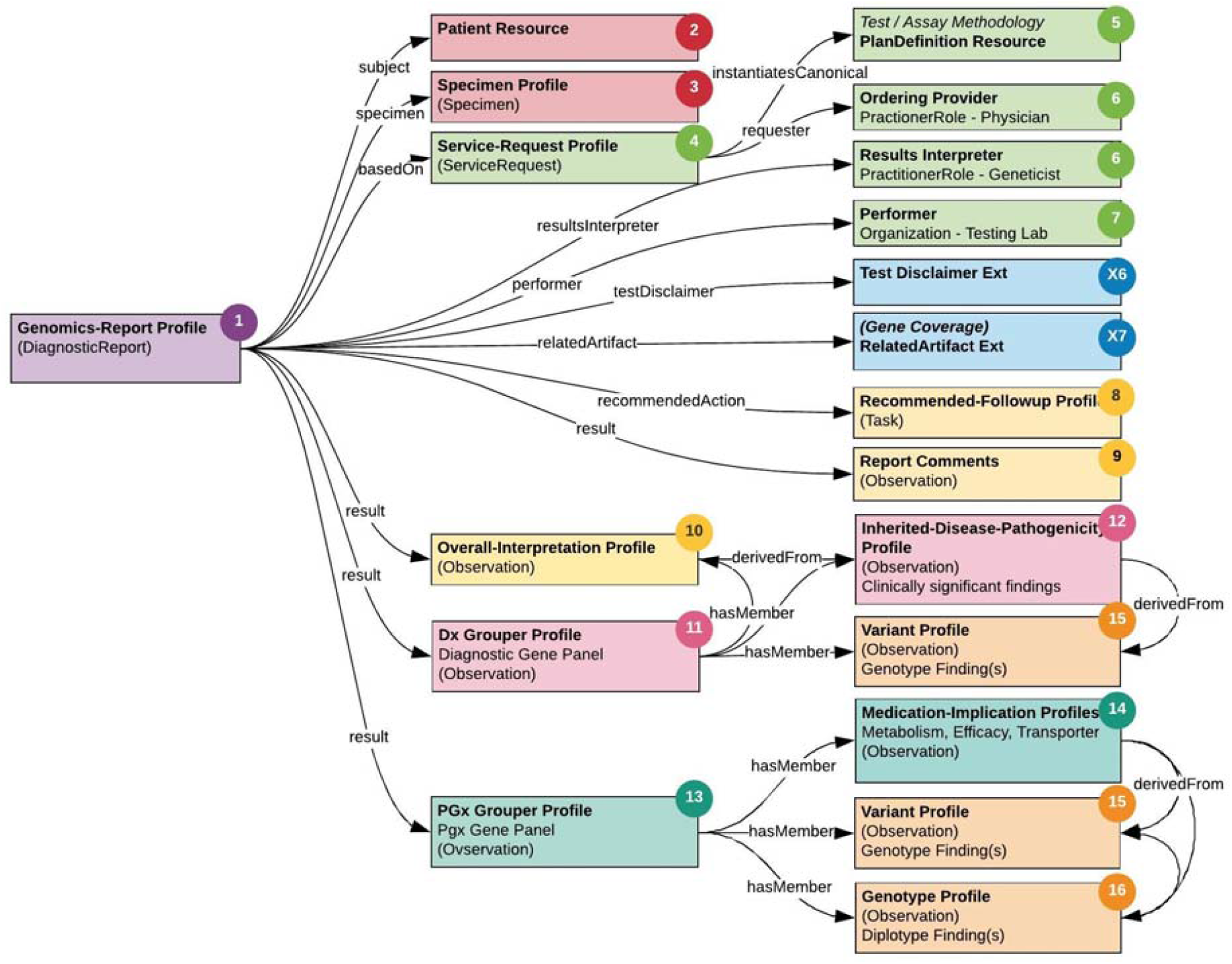
Structural Design of the eMERGE FHIR Specification. A color-coded organizational mapping of the FHIR Genomics Reporting Implementation Guide resources and profiles for eMERGE, including custom eMERGE extensions in blue (X6, X7). The color and numbering maps back to the report layout and example in Figure 4 for easier reference.

##### 3.1.2.2. Mapping eMERGE Field Level Report Elements to FHIR Resource Elements

Step 2 involved a granular mapping of every eMERGE report attribute to an equivalent field in the FHIR resources identified in the previous step. This was a laborious process which in addition to requiring precise and careful mapping of the fields themselves, also required determining naming systems and assignment of coding systems, codes and values. Online documentation[41] includes the complete set of eMERGE FHIR resources and its associated elements, with a summary listed in Table 3. Furthermore, gap analysis at this step revealed the need for additional fields such as summary interpretation text, test disclaimer etc. that were not available in the GR IG. Though we documented these as feature requests in HL7’s Jira[24], to satisfy the immediate needs of the project, we created these fields as FHIR Extensions. The full list of Extensions[42] is available on the eMERGE Read the Docs website[40].

**Table 3:**
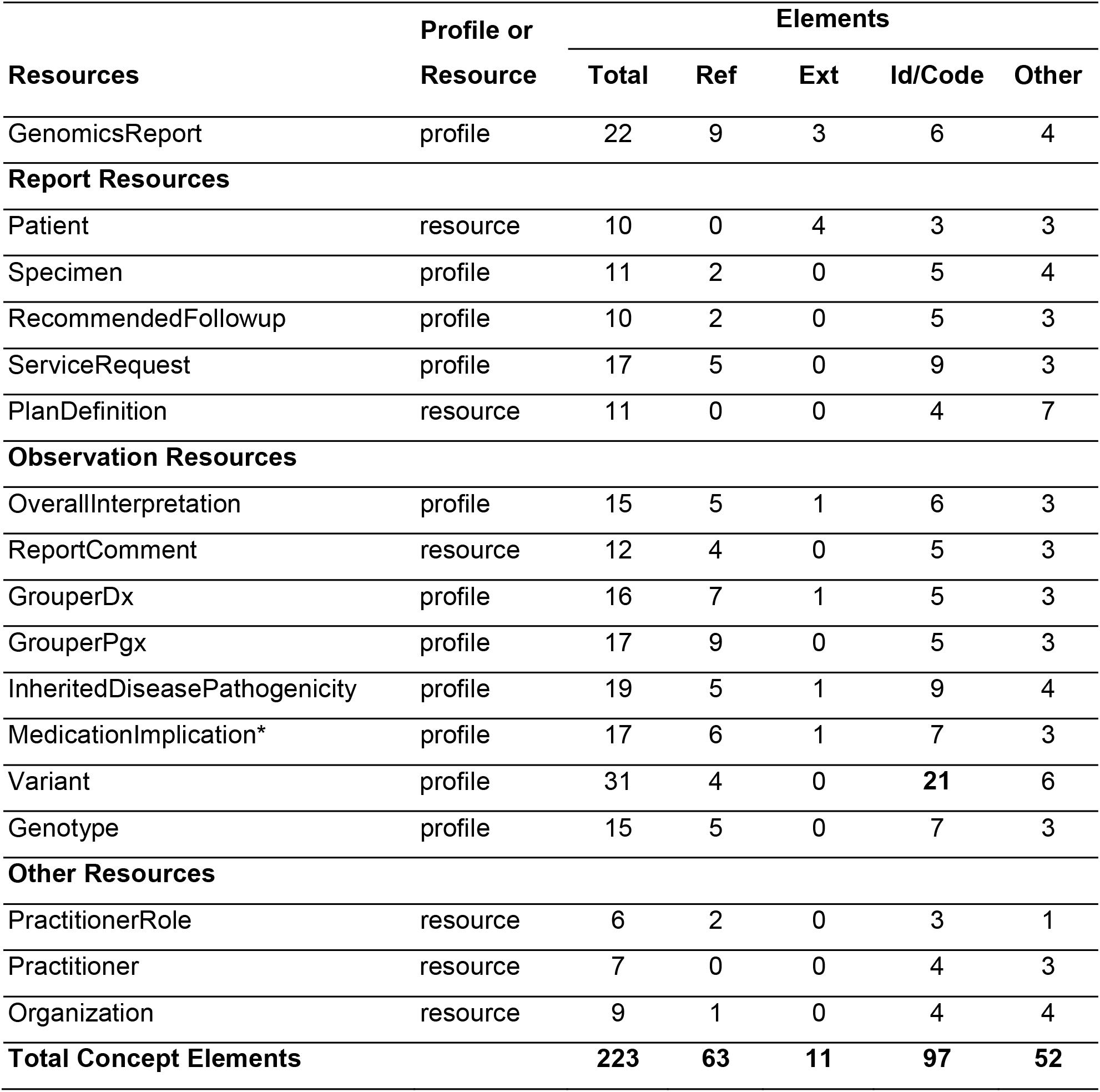
eMERGE FHIR resources. This table lists all resources and profiles in the eMERGE FHIR report bundle, with counts of the total number of elements (Total Elements) in each resource. Also included are a breakdown of the counts for each type of element i.e. references to other resources within a resource (Ref Elements), extensions used and/or created specifically for eMERGE (Ext Elements), naming or coding Systems (Id/Code Elements) and others (Other Elements).

#### 3.1.3. Harmonization and Documentation

The eMERGE FHIR Specification project was initiated while the HL7 CG WG was still working on its first draft of GR IG. Though the draft was a good start for the eMERGE Specification, it did not encompass all the eMERGE use cases. Additional documentation for the eMERGE FHIR Specification is available on the eMERGE Read the Docs website[40]; the sample reports generated via an implementation of the specification are available on GitHub[43].

### 3.2. Use Case Pilot Projects Development

The principal goals of the pilot were to a) demonstrate the successful generation of sample reports adhering to the format of the eMERGE FHIR Specification, b) successfully ingest these reports to FHIR servers at the recipient sites, and c) demonstrate use of the FHIR formatted data for NU and JHU’s respective use cases. Figure 6 illustrates the software components and data flow and is described in more detail in the following sections.

**Figure 6:**
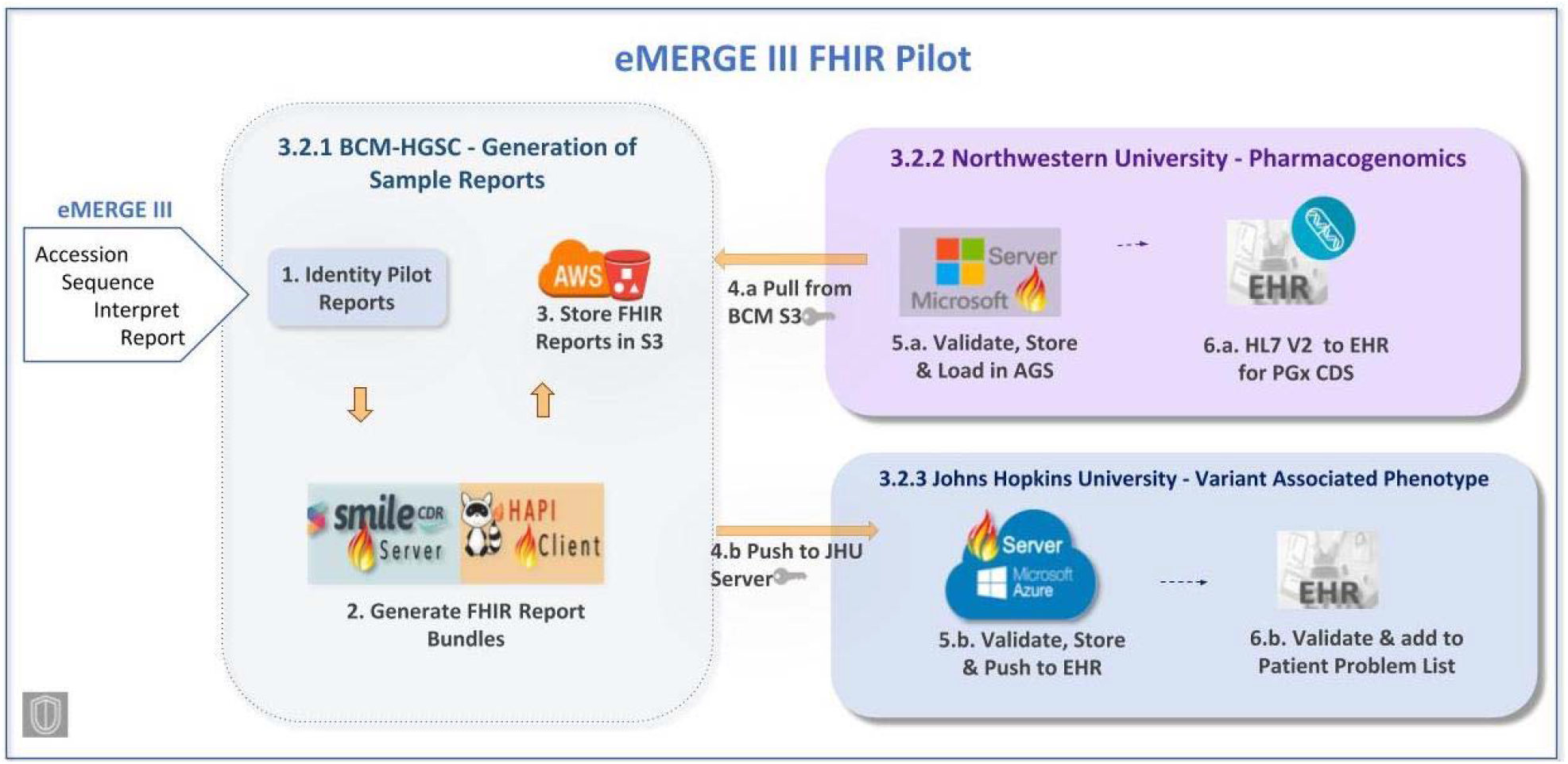
eMERGE III FHIR Pilot. FHIR bundles for the eMERGE pilot samples were generated by BCM-HGSC and pushed to NU and JHU’s FHIR servers. Data was extracted from these FHIR reports for the NU PGx and JHU Variant Associated Phenotype use cases respectively.

#### 3.2.1. BCM-HGSC - Generation of Sample Reports

##### 3.2.1.1. Pilot Sample Selection

In order to facilitate the use cases for the pilot implementation, we selected 194 sample reports from the NU eMERGE study site (161 positive and 33 negative), and included a cross section of positives, negatives, heterogeneous indications for testing, multiple variants and PGx findings, selected phenotype susceptibilities, sex and race.

Taking into account JHU’s role as a non-clinical eMERGE affiliate (meaning no genotyped data for JHU patients were available), we utilized the NU pilot set for JHU after de-identification. De-identification was performed within BCM-HGSC’s HIPAA compliant environment on DNAnexus, and according to HIPAA Safe Harbor[44]. All identifiers were stripped of their original values and left empty, with the exception of patient and sample identifiers which were replaced with randomly generated UUIDs. This follows the data security and sharing guidelines established by eMERGE Consortium’s Data Use Agreement (DUA)[45], to which all three parties involved with the pilot have agreed to comply for the pilot implementation. JHU, in their HIPAA compliant test environment, mapped these random UUIDs to patient MRNs, after matching on age, sex, and variant associated phenotypes reported, thereby creating a comparable equivalency for testing purposes.

##### 3.2.1.2. Report Generation

A total of 194 reports were successfully generated and represented as FHIR bundles. Two copies of the reports were created - one containing the original identifiers for NU, and one containing de-identified reports for JHU. No issues were found in the validation of the reports done by the study team before transmitting to NU and JHU.

A set of five report bundles, two negatives and three positives, generated with this implementation and employing the eMERGE FHIR Specification are available for download on GitHub[43]. These sample reports were generated with simulated data being used for identifiers, namespaces and results.

##### 3.1.2.3. Report Delivery

Dual modes of report delivery were employed: NU’s FHIR bundles were downloaded directly from the BCM-HGSC provisioned AWS S3 and uploaded to an open-source Microsoft FHIR Server. JHU’s FHIR bundles were pushed directly by the BCM-HGSC’s FHIRClient to an Azure Cloud Microsoft FHIR Server after successful authentication and authorization. Methods to integrate FHIR servers into the EHR at the pilot sites were considered out of scope for this paper.

#### 3.2.2. Northwestern University - Pharmacogenomics

All of the 194 FHIR bundles selected for the pilot test were loaded to the FHIR server without error. The loading process to transmit the FHIR data into the simulated AGS resulted in the creation of 194 distinct patient records, and 1358 total diplotype records (7 diplotypes per patient). All data elements needed to trigger pre-existing CDS were present in the FHIR bundles. The manual review of 25 randomly selected patients confirmed that all diplotypes were loaded without error and attributed to the correct patient. Although no patient data was transmitted to the EHR, the process resulted in the exact same data available in the NU production environment, which is known to be sufficient for triggering CDS.

#### 3.2.3. Johns Hopkins University - Variant Associated Phenotype

All of the 194 FHIR bundles selected for the pilot test were loaded to the JHU AGS without error, and the custom Validator tool verified that all 194 patients, their corresponding Specimens and all other attributes submitted by BCM-HGSC were identified (see **Figure A**.**1** in **Appendix A**). Processing the variant associated phenotype portion of the FHIR data captured in the JHU pilot AGS using the Validator tool found no discrepancies with what was expected (see **Figure A**.**2** in **Appendix A**).

#### 3.2.4. Results from Assessing Data Quality for Use Cases

Findings from analyzing data quality characteristics of the eMERGE FHIR genetic report data for the NU and JHU CDS use cases indicated high ratings for all characteristics (Completeness, Accessibility, Reliability/Accuracy, and Relevance/Fitness) with one exception (see **Table B**.**2** in **Appendix B**). The *Completion* characteristic was rated lower for the JHU use case due to a need for additional processing of the report data in order to add a variant associated phenotype to a patient problem list using the FHIR API. Results are described in more detail in **Appendix B**.

## 4. Discussion

Through this eMERGE pilot effort we developed a standards-based specification for representing clinical genetic and genomic test results and demonstrated its viability with EHR integration and clinical use cases. This project supported the development of the FHIR Specification for genomics by basing the eMERGE FHIR Specification on the GR IG, and subsequently demonstrating use of the FHIR standard for genomic medicine. Through a data quality assessment framework (**Appendix B**), we confirmed that the data structures of the FHIR-formatted reports were acceptable for the two CDS use cases. Additionally, the deployment provided several lessons that guide future evolution of the FHIR Specification.

The CG WG released the first standard trial use (STU1) of the GR IG in November 2019, nearly a year after the work described in this manuscript began. As an early adopter, we worked collaboratively with the CG WG to further inform the design of the STU1 specification as it matured. As our experience demonstrated, feedback from early adopters contributes significantly to the development of a draft specification, but adopters must be flexible as the standard evolves in parallel.

The level of detail in the GR IG, which uses a series of FHIR profiles designed to reflect the complexity of clinical genomic test results, requires adopters to have an in-depth understanding of the domain as well as time and resources to implement the complex specification itself. This could be a deterrent to adoption for vendors, laboratories, or consumers that have relatively simple data.

The CG WG, recognizing that the complexity of the GR IG could be a barrier to adoption, is actively working with stakeholders to refine the specification. The experience of this project demonstrates the potential value of working directly with adopters to identify minimum viable products that are based on local use cases. Reciprocally, the CG WG would benefit significantly from concerted efforts by potential adopters (i.e. vendors, laboratories, healthcare systems, health IT engineers) through participation and collaboration to help inform the roadmap and development of the GR IG. Examples of such initiatives include ONC Sync for Genes[46], PGx CDS with FHIR and CDS Hooks[47], HLA Reporting with FHIR[48], Variant Interpretation for the Cancer Consortium[49], and mCODE[50].

### 4.4. Future Considerations

The issues we addressed during this project varied in scale in both importance as well as complexity. From these we have selected four key possibilities for further development: variant data types, interpretation summary text, gene/region coverage, and PGx results representation. A comprehensive description of all the issues can be found in the eMERGE FHIR Specification[36].

#### 4.4.1. Variant Data Types

Community standards are required to define variants in a manner that will allow science and medicine to adopt and develop tools and procedures to track and attach information to variants at the scale demanded by genetic testing. Efforts to describe and identify variants include: Human Genome Variation Society (HGVS) nomenclature[51], International System for Human Cytogenetic Nomenclature (ISCN) nomenclature[52], and Variant Call File (VCF) formats[53]. Public repositories and registries established by various authorities have evolved over the past decade or more to provide identifiers for variants in these systems (e.g. dbSNP[54], ClinVar[55], ClinGen Allele Registry[56], COSMIC[57]).

The notion of a definitional or reference variant does not exist in FHIR or the GR IG, with FHIR and the CG WG lacking the data types and resources needed to build these referenceable variants. The CG WG’s proposed resolution is to develop FHIR profiles based on the Observation resource. This begins to address the concern of separating the variant observation from the other assertion profiles, like disease causality or therapeutic implications of metabolism or efficacy. However, it does not provide a pragmatic computational standard for representing the variants referenced within those observed variants and associated assertions.

The Genomic Knowledge Standards (GKS) Workstream of the Global Alliance for Genomic Health (GA4GH) is developing and expanding the Variation Representation Specification[58,59] (VRS) to address the need for standards for computationally sharing variation. Instituting such a model in FHIR will significantly reduce the adoption risks caused by the complexity and unguided extensibility of the current GR IG and FHIR Specifications. As such, the growing collaboration between the CG WG and the GA4GH GKS Workstream represents a promising step forward at introducing the concepts, resources and data types needed in the FHIR Specification to improve the viability of implementing use cases related to variation in FHIR systems.

#### 4.4.2. Interpretation Summary Text

While structured and coded results are of great importance to the computational utility of results, unstructured text will always play a significant role in conveying information between humans. Though there are a number of text attributes available throughout the GR IG resources, the genetics community requires the ability to associate an interpretation summary with the overall results of the report and with one or more classified variants. It is our recommendation that the CG WG consider all of the important kinds of text fields needed to support clinical genetic test results and assure that there is a mechanism to do so, starting with an interpretation summary text field.

#### 4.4.3. Gene / Region Coverage

Because testing methods vary, it is important to provide a quantitative representation of the precise molecular sequenced regions covered and the quality of coverage for each region. Perhaps more importantly, this information clearly identifies which regions of the genome were not covered but may be relevant for the indication for testing. Receiving systems would then be able to accurately determine whether a patient may need follow-up testing to interrogate additional regions, and results across cohorts and studies would be more comparable.

#### 4.4.4. PGx Results Representation

Key challenges to computationally representing PGx results are 1) to include the assayed variants that underlie the derived haplotype/diplotype assertions and 2) to maintain a clear distinction between the case-independent PGx knowledge statements about the haplotype/diplotype states and the patient-specific PGx implications expressed in the interpretation, which are based on the assay findings. The GR IG should provide a mechanism for defining variants at various levels that can be referenced either in the context of a patient observation or independently in a generalizable knowledge base. Also, it is currently difficult to convey the underlying assayed variants using the existing specification; if this could be accomplished then clinical systems could identify previously reported PGx results that may be impacted by future knowledge updates. This design approach supports the need for variant data types as discussed in section 4.4.1.

## 5. Conclusion

With this project, we have developed and demonstrated the use of a structured, computable and interoperable HL7 FHIR standard for eMERGE Clinical Reporting and have demonstrated the feasibility of implementing a standardized representation of clinical genetic and genomic results in the EHR. We have helped move the needle forward for the HL7 Clinical Genomics Workgroup in its development of the Genomics Reporting Implementation Guide by identifying and contributing to several growth areas[36] in order to effectively fulfill the return of clinical genomics results.

The two pilot use cases helped us understand the landscape to use FHIR-formatted reports for genomic medicine and its challenges. Establishing a standard, though fundamental, is but one step towards the effective use of genomic medicine in the clinic. In order to achieve this vision, additional work is needed not only to mature and stabilize a standard for structured representation of clinical genomic data but also with establishing engagement and collaboration between domain experts. Myriad perspectives are needed, including that of clinicians and geneticists, standards bodies such as HL7, EHR and other clinical system vendors, diagnostic laboratories, and health IT. These perspectives will further identify genomic medicine use cases and build standards and systems to support these use cases as well as target integration and interoperability amongst users, generators and consumers of clinical genomic data. Tenacious continuation of such undertakings and joint complementary efforts is the path towards supporting standardization and effective use of genomic data in the clinic.

## Acknowledgements

1. Members of the HL7 Clinical Genomics Workgroup;
2. Members of the eMERGE EHRI Workgroup;
3. Steve Ordhal contributed through Microsoft’s Partner Network that provides enhanced services to Johns Hopkins University.
4. Members of the BCM-HGSC Clinical Laboratory

## Funding

This work was conducted under Phase III of the eMERGE Network, which was initiated and funded by the NHGRI through the following grants: U01HG008657 (Kaiser Permanente Washington/University of Washington); U01HG008685 (Brigham and Women’s Hospital); U01HG008672 (Vanderbilt University Medical Center); U01HG008666 (Cincinnati Children’s Hospital Medical Center); U01HG006379 (Mayo Clinic); U01HG008679 (Geisinger Clinic); U01HG008680 (Columbia University Health Sciences); U01HG008684 (Children’s Hospital of Philadelphia); U01HG008673 (Northwestern University); U01HG008701 (Vanderbilt University Medical Center serving as the Coordinating Center); U01HG008676 (Partners Healthcare/Broad Institute); U01HG008664 (Baylor College of Medicine); and U54MD007593 (Meharry Medical College).

## Notes

### Competing Interest Statement

Luke V. Rasmussen has a patent GENERATING DATA IN STANDARDIZED FORMATS AND PROVIDING RECOMMENDATIONS that is no longer being pursued.
Samuel J. Aronson, Hana Zouk and Heidi L. Rehm are employed by Mass General Brigham which receives royalties on sales of GeneInsight software.
David R. Crosslin is a consultant for UnitedHealth Group.
Richard A. Gibbs declares that Baylor College of Medicine receives payments from Baylor Genetics Laboratories, which provides services for genetic testing; Baylor College of Medicine is part owner of Codified Genomics.
Eric Venner is a cofounder of Codified Genomics, which provides variant interpretation services.
All other authors declare no competing interests.

