## Appendix A for "Genomic Considerations for FHIR; eMERGE Implementation Lessons"

### Appendix A - Johns Hopkins University Technical Validation

***Account access validation:*** To validate the BCM-HGSC and JHU service account access to the JHU AGS we used Postman[[1]](https://www.zotero.org/google-docs/?sVsK16) to perform the following two steps : (1) authenticate the account to MS Azure Active Directory (AAD) and obtain a bearer token; and (2) use the token to perform an action. To perform step 1, we did a POST to the AAD using a client id, a client secret and a tenant id. The expected result was an expiring token for the BCM-HCSC and JHU accounts. To perform step 2, we did a GET with the tokens to see if the accounts could read data and a POST with the tokens to see if the account could write data. The BCM-HGSC service account’s role was solely to send data (i.e., POST to the FHIR server), and the JHU service account should be able to both read and write data (i.e., GET and POST). The expected results for POST and GET actions were to complete with no error responses. Figure A.1 illustrates what an error response would look like due to an attempt to read data using the BCM-HGSC (write-only) service.

***Import process validation:*** The BCM-HGSC created an application to create FHIR transaction bundles and submit them to the JHU AGS, and the JHU team wrote a Validator tool parse the BCM-HGSC log file and get the submitted data from the AGS. That BCM-HGSC application logged its submission to a log file, in the order that the application builds the bundles and sends them. The Validator tool then parsed the BCM-HGSC log file and retrieved the submitted data from the AGS. As the JHU Validator tool parsed the log file, it kept a count of each of the different FHIR types submitted. The tool then retrieved each of the submissions it found in the log file. The submitted count was then compared to the count it found in the JHU AGS. In addition, the variant associated phenotype portion of the FHIR data was checked to verify that the count of OMIM identifiers in the reports matched what was expected. Figure A.2 illustrates the output of the custom Validator tool indicating that all 194 patients, their corresponding Specimens, and all other attributes submitted by BCM-HGSC. Figure A.3 illustrates the output of the Validator tool indicating that the count of OMIM identifiers across all submissions matched what was expected.

**Figure A.1 Authorization response**. The figure displays the response when the BCM-HGSC service account attempts to do a GET.

{

"resourceType": "OperationOutcome",

"id": "1cb35ca6-a88d-426d-aa23-cc11f12f83be",

"issue": [

{

"severity": "error",

"code": "auth-access",

"diagnostics": "User/Application must be in a reader role to access"

}

]

}

**Figure A.2 Abbreviated BCM-HGSC submission validation results**.

2020-05-06 10:24:08 Validator:109 - All Patient objects accounted for 194 : 194

2020-05-06 10:25:18 Validator:109 - All Specimen objects accounted for 194 : 194

2020-05-06 10:26:07 Validator:109 - All ServiceRequest objects accounted for 194 : 194

2020-05-06 10:41:29 Validator:109 - All Observation objects accounted for 3632 : 3632

2020-05-06 10:44:38 Validator:109 - All Practitioner objects accounted for 776 : 776

2020-05-06 10:46:13 Validator:109 - All Organization objects accounted for 388 : 388

2020-05-06 10:47:44 Validator:109 - All PractitionerRole objects accounted for 388 : 388

2020-05-06 10:48:33 Validator:109 - All PlanDefinition objects accounted for 194 : 194

2020-05-06 10:49:19 Validator:109 - All DiagnosticReport objects accounted for 194 : 194

**Figure A.3 Abbreviated OMIM subject validation results**.

2020-07-09 12:29:46 OmimCheck:110 - OMIM identifier: 114480 Subject Count: 10

2020-07-09 12:29:46 OmimCheck:110 - OMIM identifier: 120435 Subject Count: 2

2020-07-09 12:29:50 OmimCheck:110 - OMIM identifier: 176807 Subject Count: 10

2020-07-09 12:29:53 OmimCheck:110 - OMIM identifier: 208900 Subject Count: 10

2020-07-09 12:29:56 OmimCheck:110 - OMIM identifier: 604370 Subject Count: 10

2020-07-09 12:29:57 OmimCheck:110 - OMIM identifier: 609310 Subject Count: 3

2020-07-09 12:30:00 OmimCheck:110 - OMIM identifier: 612555 Subject Count: 10

2020-07-09 12:30:01 OmimCheck:110 - OMIM identifier: 613348 Subject Count: 2

2020-07-09 12:30:04 OmimCheck:110 - OMIM identifier: 614337 Subject Count: 3

2020-07-09 12:30:05 OmimCheck:110 - OMIM identifier: 614350 Subject Count: 4
