## Appendix B for "Genomic Considerations for FHIR; eMERGE Implementation Lessons"

### Appendix B - Data Quality Assessment

***Methods:*** Our approach to assess the data quality characteristics of the eMERGE FHIR genetic report data used by the NU and JHU for their CDS use cases, adapted a scoring system[[2]](https://www.zotero.org/google-docs/?KKuFwx) which scores four characteristics (Completeness, Accessibility, Relevance/Fitness, and Reliability/Accuracy) on a scale of 0-2, which is shown in **Table B.1**. It was applied by NU and JHU following the completion of their use case, and the scores were discussed collectively to provide additional rationale and context.

***Results:*** The results from assessing data quality characteristics for the NU and JHU use cases are summarized in **Table B.2**. For the NU use case, content and format validity scores of 2 were assigned for all four assessed characteristics. The *Completeness* characteristic was rated as a 2 because all necessary data elements were available to trigger the pre-existing PGx CDS rules, and *Accessibility* was rated as a 2 because all data elements could be accessed without issue. For the *Reliability/Accuracy* category, the manual review of 25 randomly selected reports confirmed that all diplotypes were loaded correctly and attributed to the correct patient. Finally, the *Relevance/Fitness* characteristic was also rated as a 2 since the data available and medical vocabularies used were usable without additional transformation.

For the JHU use case, a score of 1 was assigned for the *Completion* characteristic because we found that in order to use the FHIR API to add a variant associated phenotype to a patient problem list, some additional processing was required. In particular, condition annotations needed to be created and added as Condition resources to the FHIR reports. A score of 2 was assigned for all other characteristics. For the *Accessibility* characteristic, the FHIR API provided the data access we needed. For the *Relevance/Fitness* category, upon adding a condition to a problem list using the FHIR API, it is added as “unverified” by default and provides clinicians an opportunity to verify, reject, or change a problem to something different prior to adding it to the patient problem list. For the *Reliability/Accuracy* category, our assessment of variant associated phenotypes found no discrepancies with what was expected and verified that they were loaded correctly.

| **Characteristic** | **Description and framing question** | **Scores** |
| --- | --- | --- |
| Completeness | Data were considered missing when they did not exist in an electronic form.  “There exists a deficiency of a component that will impact use of the data” | - No-using eMERGE report FHIR resources only (2) - No-using FHIR resources outside of those in the eMERGE reports (1) - Yes (0) |
| Accessibility | Data were considered inaccessible when they existed in an electronic form but could not be easily retrieved or used.  “A data access interface is provided” | - Yes-using FHIR API (2) - Yes-custom API(1) - No-not possible (0) |
| Relevance/Fitness | Ability to adapt to individual and clinical contexts.  “Degree to which the data produced matches user’s needs” | - Completely (2) - Partially (1) - Not at all (0) |
| Reliability/Accuracy | Accuracy of a given data value is determined by comparing it to a known reference value.  “Data representation (or value) reflects the true state of the source information” | - Completely (2) - Partially (1) - Not at all (0) |

**Table B.1:** Assessment of the characteristics of the eMERGE FHIR genetic report data used by the NU and JHU for their CDS use cases using a data quality scoring system[[2]](https://www.zotero.org/google-docs/?qx0xOf).

|  | **NU PGx CDS** | **JHU Variant Associated Phenotype CDS** |
| --- | --- | --- |
| Completeness | 2-no-using eMERGE report  FHIR resources only | 1-no-using FHIR resources outside of those in the eMERGE reports |
| Accessibility | 2-yes-using FHIR API | 2-yes-using FHIR API |
| Relevance/Fitness | 2-completely | 2-partially |
| Reliability/Accuracy | 2-completely | 2-completely |

**Table B.2:** Summary of data quality characteristics of the eMERGE FHIR genetic report data used by the NU and JHU for their CDS use cases.
